# ATR inhibition augments the efficacy of lurbinectedin in small cell lung cancer

**DOI:** 10.1101/2022.12.16.520783

**Authors:** Christopher W. Schultz, Yang Zhang, Rajaa Elmeskini, Astrid Zimmermann, Haiqing Fu, Yasuhisa Murai, Darawalee Wangsa, Suresh Kumar, Nobuyuki Takahashi, Devon Atkinson, Liton Kumar Saha, Chien-Fei Lee, Brian Elenbaas, Parth Desai, Robin Sebastian, Thomas Ried, Mirit Aladjem, Frank T. Zenke, Zoe Weaver, Yves Pommier, Anish Thomas

## Abstract

Small cell lung cancer (SCLC) is the most lethal type of lung cancer. Specifically, MYC-driven non-neuroendocrine SCLC are particularly resistant to standard therapies. Lurbinectedin was recently approved for the treatment of relapsed SCLC, but combinatorial approaches are needed to increase the depth and duration of responses to lurbinectedin. Using high-throughput screens, we found inhibitors of ataxia telangiectasia–mutated and rad3-related (ATR) as the most effective agents for augmenting lurbinectedin efficacy. First-in-class ATR inhibitor berzosertib synergized with lurbinectedin in multiple SCLC cell lines, organoid and *in-vivo* models. Mechanistically, ATR inhibition abrogated S-phase arrest induced by lurbinectedin and forced cell-cycle progression causing mitotic catastrophe and cell death. High *CDKN1A*/p21 expression was associated with decreased synergy due to G1 arrest, while increased levels of *ERCC5*/XPG were predictive of increased combination efficacy. Importantly, MYC-driven non-neuroendocrine tumors which were resistant to first-line therapies displayed decreased *CDKN1A*/p21 expression and increased *ERCC5*/XPG indicating they were primed for response to lurbinectedin-berzosertib combination. The combination is being assessed in a clinical trial NCT04802174.

**Graphical Abstract:** 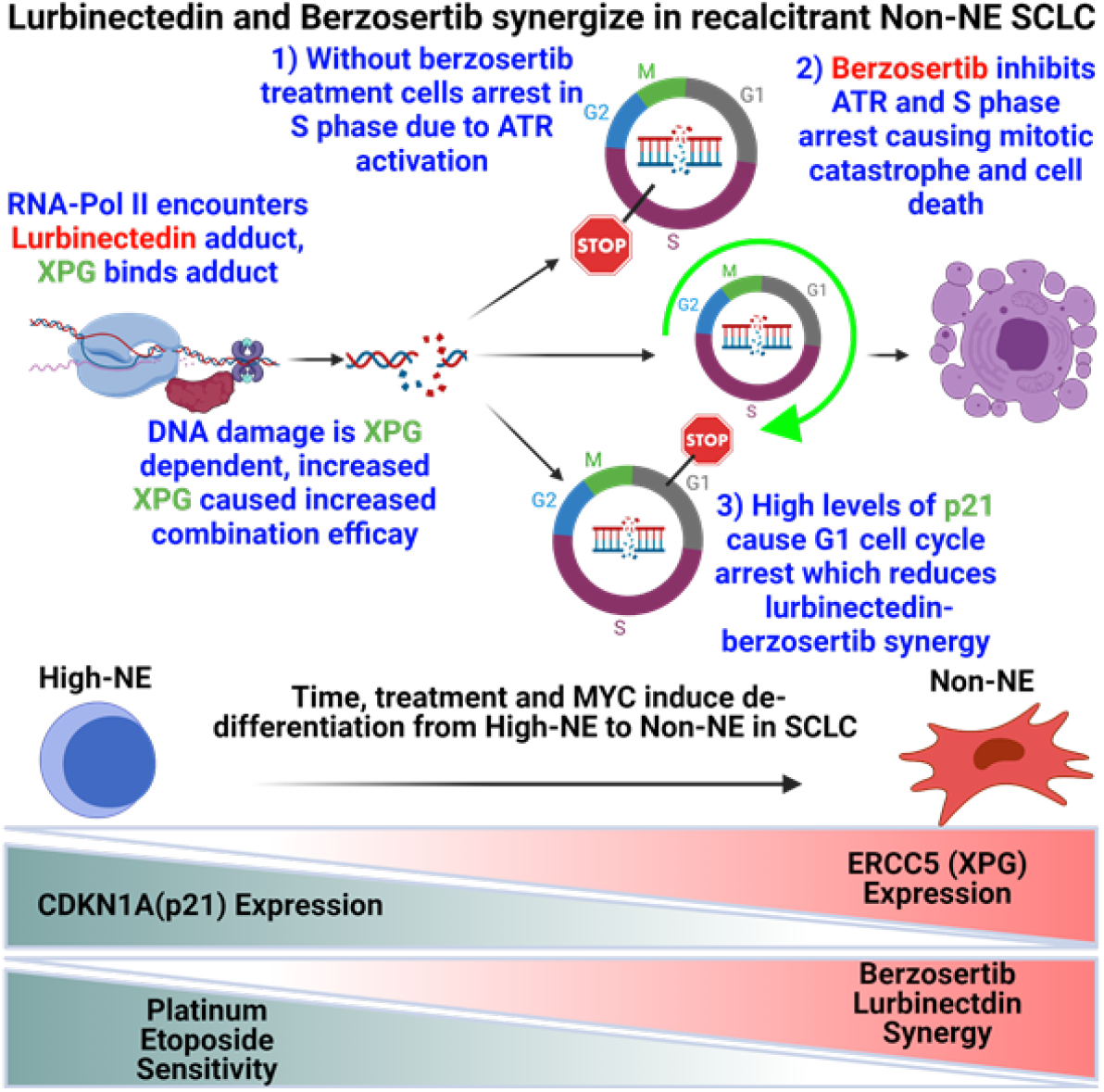

## Introduction

Despite major advances in targeting oncogenes and the immune inhibitory checkpoints, most cancer patients die from chemotherapy-resistant disease. Multiple resistance mechanisms to chemotherapy have been described. For DNA damaging chemotherapies, reduced intracellular drug intake, intracellular inactivation of the agent, increased DNA repair, activation of alternative DNA repair pathways, and impaired apoptotic signaling are among the most common resistance mechanisms[1].

Targeting DNA repair pathways using combination strategies represents a rational approach to overcome chemotherapy resistance. The ataxia telangiectasia–mutated and rad3-related (ATR) kinase is a master regulator of DNA damage response that plays a key role in stabilizing the genome when DNA replication is compromised [2]. ATR is activated by regions of single-stranded DNA (ssDNA), commonly generated as a result of DNA replication stress produced pharmacologically or by oncogene activation [3]. Once activated, ATR functions to safeguard genomic integrity and safeguard replication by slowing the progression of replication forks, inhibiting distal replication origin firing, ensuring sufficient supply of deoxynucleotides, and promoting cell cycle arrest primarily by activation of intra S and G2/M cell cycle checkpoints [3]. Accordingly, ATR inhibition leads to the loss of the S and G2/M checkpoints, allowing cells with damaged DNA to progress prematurely into M phase, leading to mitotic catastrophe and cell death [3, 4]. As such, multiple potent and selective ATR inhibitors are in pre-clinical and clinical development for cancer therapy [4-8].

SCLC is a neuroendocrine (NE) tumor characterized by near universal bi-allelic loss of tumor suppressors *TP53* and *RB1*. In addition, amplification of *MYC* family genes occurs in ∼20% of SCLC and overexpression in ∼50% of SCLC [9, 10]. Most patients are diagnosed with widely metastatic disease, and are treated with a combination regimen of platinum, etoposide, and immunotherapy. Despite initial responses, risk of relapse is high with >90% of patients progressing within two years [11]. Second-line treatment options include topotecan and lurbinectedin, but the depth and duration of responses are modest and most relapsed tumors do not respond to additional chemotherapy. Furthermore, SCLCs have few targetable alterations [9] and tend not respond to therapies targeted at somatic mutations [12]. Recent studies have revealed heterogeneity of the SCLC neuroendocrine cell state, with tumors consisting of cells with NE and non-neuroendocrine (non-NE) features [13]. SCLC heterogeneity increases over the course of treatment, with an increase in chemo-resistant non-NE cells evolving over time [14]. Importantly, NE differentiation is emerging as a potential predictor of response to therapy [6, 15-17].

Lurbinectedin is a synthetic alkylating derivative of trabectedin that was recently approved for SCLC patients with disease progression on or after platinum-based chemotherapy [18]. Lurbinectedin covalently binds to DNA forming adducts that irreversibly stall elongating RNA polymerase II (Pol II) on the DNA template, generating DNA double-strand breaks (DSBs) [19, 20]. Here we report an exquisite, NE differentiation-dependent synergistic interaction between the ATR inhibitor berzosertib and lurbinectedin. Mechanistically, lurbinectedin induces DSBs and activates ATR causing cell cycle arrest allowing for repair and resistance, this process is inhibited with berzosertib co-treatment. This combination produced greater synergy in MYC-high, chemo-resistant, non-NE SCLC subtype. Notably, these findings form the basis for a clinically actionable combination of a DNA repair inhibitor and a DNA-damaging agent to overcome SCLC chemoresistance.

## Materials and Methods

### Cell lines

NCI-H211, NCI-H524, DMS-114, NCI-H841, NCI-H1048, NCI-H1341, NCI-H446, NCI-H146, NCI-H889 and U2OS DRGFP [21] cell lines were purchased from ATCC. NCI-H211, NCI-H889, NCI-H1048, NCI-H1341 and U2OS DRGFP cell lines are female and the rest are male, additional information for cell lines can be observed in Table 1. Cell lines were authenticated using short tandem repeat analysis, and were monthly tested for mycoplasma contamination. Cell media was RPMI-1640 supplemented with 10% FBS for all lines to maintain consistency. DT40 (chicken cell lines) were grown in Dulbecco”s Modified Eagle”s Medium (DMEM) supplemented with FBS 10% and chicken serum 5%. Cells were grown at 37°C and 5% CO2.

**Table 1.**
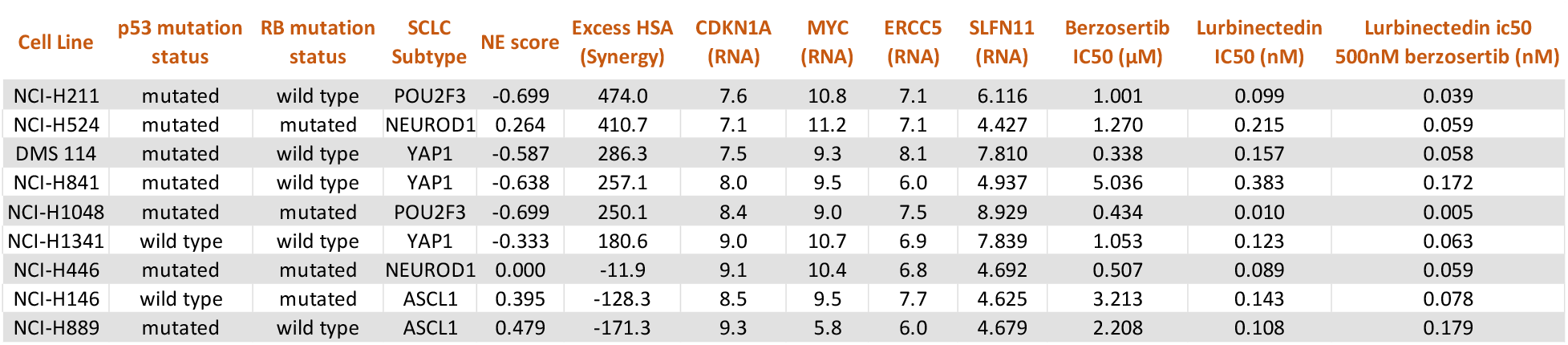
Critical variables for lurbinectedin and berzosertib efficacy and synergy in SCLC cell lines: Pertinent values from the 9 SCLC cell lines utilized in this work. All RNA values are given as log 2 values and these along with NE scores, RB (RB1 and RB2 both assessed, all mutations in RB1) and p53 mutation status are publicly available at https://discover.nci.nih.gov/rsconnect/cellminercdb.

| Cell Line | p53 mutation status | RB mutation status | SCLC Subtype | NE score | Excess HSA (Synergy) | CDKN1A (RNA) | MYC (RNA) | ERCC5 (RNA) | SLFN11 (RNA) | Berzosertib IC50 (μM) | Lurbinectedin IC50 (nM) | Lurbinectedin ic50 500nM berzosertib (nM) |
| --- | --- | --- | --- | --- | --- | --- | --- | --- | --- | --- | --- | --- |
| NCI-H211 | mutated | wild type | POU2F3 | -0.699 | 474.0 | 7.6 | 10.8 | 7.1 | 6.116 | 1.001 | 0.099 | 0.039 |
| NCI-H524 | mutated | mutated | NEUROD1 | 0.264 | 410.7 | 7.1 | 11.2 | 7.1 | 4.427 | 1.270 | 0.215 | 0.059 |
| DMS 114 | mutated | wild type | YAP1 | -0.587 | 286.3 | 7.5 | 9.3 | 8.1 | 7.810 | 0.338 | 0.157 | 0.058 |
| NCI-H841 | mutated | wild type | YAP1 | -0.638 | 257.1 | 8.0 | 9.5 | 6.0 | 4.937 | 5.036 | 0.383 | 0.172 |
| NCI-H1048 | mutated | mutated | POU2F3 | -0.699 | 250.1 | 8.4 | 9.0 | 7.5 | 8.929 | 0.434 | 0.010 | 0.005 |
| NCI-H1341 | wild type | wild type | YAP1 | -0.333 | 180.6 | 9.0 | 10.7 | 6.9 | 7.839 | 1.053 | 0.123 | 0.063 |
| NCI-H446 | mutated | mutated | NEUROD1 | 0.000 | -11.9 | 9.1 | 10.4 | 6.8 | 4.692 | 0.507 | 0.089 | 0.059 |
| NCI-H146 | wild type | mutated | ASCL1 | 0.395 | -128.3 | 8.5 | 9.5 | 7.7 | 4.625 | 3.213 | 0.143 | 0.078 |
| NCI-H889 | mutated | wild type | ASCL1 | 0.479 | -171.3 | 9.3 | 5.8 | 6.0 | 4.679 | 2.208 | 0.108 | 0.179 |

### Organoids

Human Small Cell Lung Cancer (SCLC) autopsies/biopsy specimens were obtained from the CCR-NCI biobank following patient consent, NIH institutional review board (IRB) and ethical approval. The pathological specimens were immediately stored in storage media (1x DMEM/F12, 1X Glutamax and 10mM HEPS buffer) on ice. The tissues were immediately subjected to enzymatic disassociation. Human SCLC organoids were cultured in Minimal Basal Media (MBM) as described previously (Ref). Briefly, PDOs were culture in drop of growth factor reduced Basement Membrane Extract (BME) (Corning) and medium was refreshed every four days. The culture media contains, DMEM/F12(Gibco) with 50ng/ml EGF (StemCell Technologies), 100nM IGF-1 (StemCell Technologies), 1x N2 Supplement (Thermofisher Scientific), 1x B27 (Themofisher Scientific) and 10 µM Y-27632(StemCell Technologies). The organoids were passage through shear stress with cold 1 U/ml Dispase/DMEM/F12 solution (StemCell Technologies) followed by Trypsin-EDTA (Invitrogen). Organoids were grown at 37°C and 5% CO2.

### Mouse tumor models

Eight-week-old male or female NSG mice (NOD.Cg-Prkdc scid Il2rg tm1Wjl/SzJ; # 005557, The Jackson Laboratory, Bar Harbor, ME) were implanted subcutaneously with fresh patient-needle biopsy supported with Matrigel (Corning) to generate our PDX model. Mice were treated weekly with lurbinectedin at 0.18mg/kg intravenously on day 1, and berzosertib at 20mg/kg intraperitoneally on day1/5. For our pharmacodynamic study mice were harvested 24h after the second dose Lurbinectidin and berzosertib. The Animal Study Protocol was approved and followed the Frederick National Laboratory Animal Care and Use Committee guidelines.

For the DMS 114 xenograft mouse model female H2dRag2 mice(C;129P2-H2d-Rag2<tm1Fwa IL2rgtm1; Taconic, Denmark), 8-10 weeks old mice were used. Mice were treated weekly with weekly cycles consisting of the following arms: vehicle, lurbinectedin alone, berzosertib alone, lurbinectedin day 1 berzosertib day 1, lurbinectedin day 1 berzosertib day 2 and lurbinectedin day 1 berzosertib days 1 and 2. Lurbinectidin was dosed at 0.18mg/kg intravenously, and berzosertib (50mg/kg oral). The study design and animal usage were approved by local animal welfare authorities (Regierungspräsidium Darmstadt, Hesse, Germany, protocol registration number DA4/Anz.1040).

### Screening and Synergy

The screen analyzed in Figure 1A,B,C and Supplemental Tables 1 and 2 was performed in NCI-H446 SCLC cells as described previously [6], the data for this screen (12831) is available at https://matrix.ncats.nih.gov/. For further synergy analysis, cells were seeded at 1,000 cells per well in 384 well plates, and collected using Cell Titer Glo (Promega, Madison, WI, USA) 72 hours after drug treatment. Highest single agent (HSA) synergy was determined by calculating the difference between the most effective single agent and the combination of agents. All matrix formats stated in this work will be presented as lurbinectedin x berzosertib with one control, i.e. a 10x6 matrix would have 6 concentrations of lurbinectedin with one being zero across 6 concentrations of berzosertib with one being zero. For thymidine based experiments cells were plated at 1,000 cells per well in 384 well plates. The next day they were treated with thymidine 2mM for 18 hours followed by a 6-9 hour release and then treated with thymidine at 2mM +/-1nM lurbinectedin +/-2µM berzosertib for 72 hours.

**Table 2.**
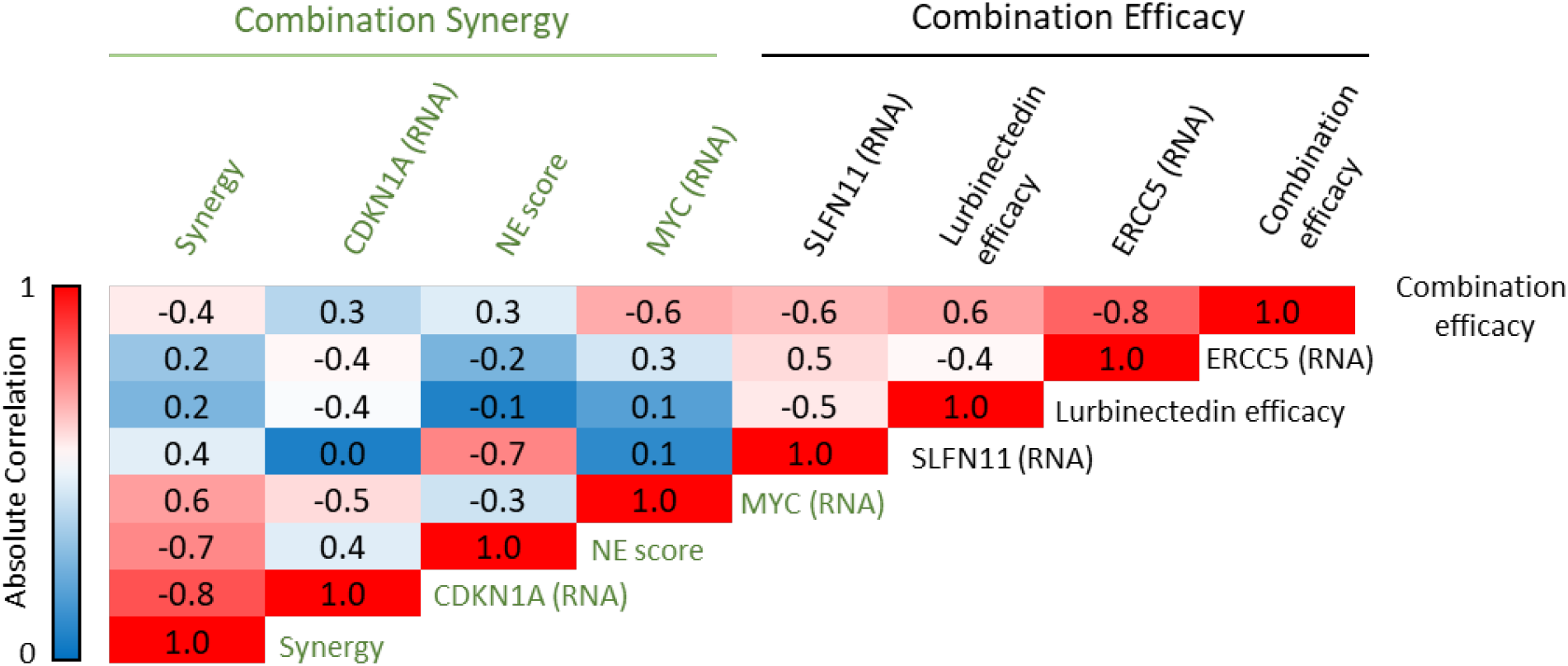
Combination and synergy determinants for the combination of lurbinectedin and berzosertib in SCLC: We have determined that synergy of lurbinectedin and berzosertib is governed by *CDKN1A* (p21) which in turn is regulated by NE differentiation and *MYC* expression. The efficacy of the combination (IC50 of lurbinectedin in the presence of 500nM berzosertib) correlates most strongly with *ERCC5* (XPG) expression followed by lurbinectedin solo agent efficacy (ic50) and *SLFN11* expression. Here the numbers indicate Pearson correlation values of variables from our panel of nine cell lines while the color scale is indicative of absolute Pearson correlation values. As synergy is denoted as HSA values and combination efficacy is representative of ic50 values of lurbinectedin in the presence of 500nM berzosertib, positive correlation with synergy indicates greater synergy while negative correlation with efficacy indicates greater efficacy (i.e. MYC RNA correlates positively with synergy and negatively with efficacy indicating with greater MYC expression there is more synergy and increased efficacy of the combination).

**Fig. 1.**
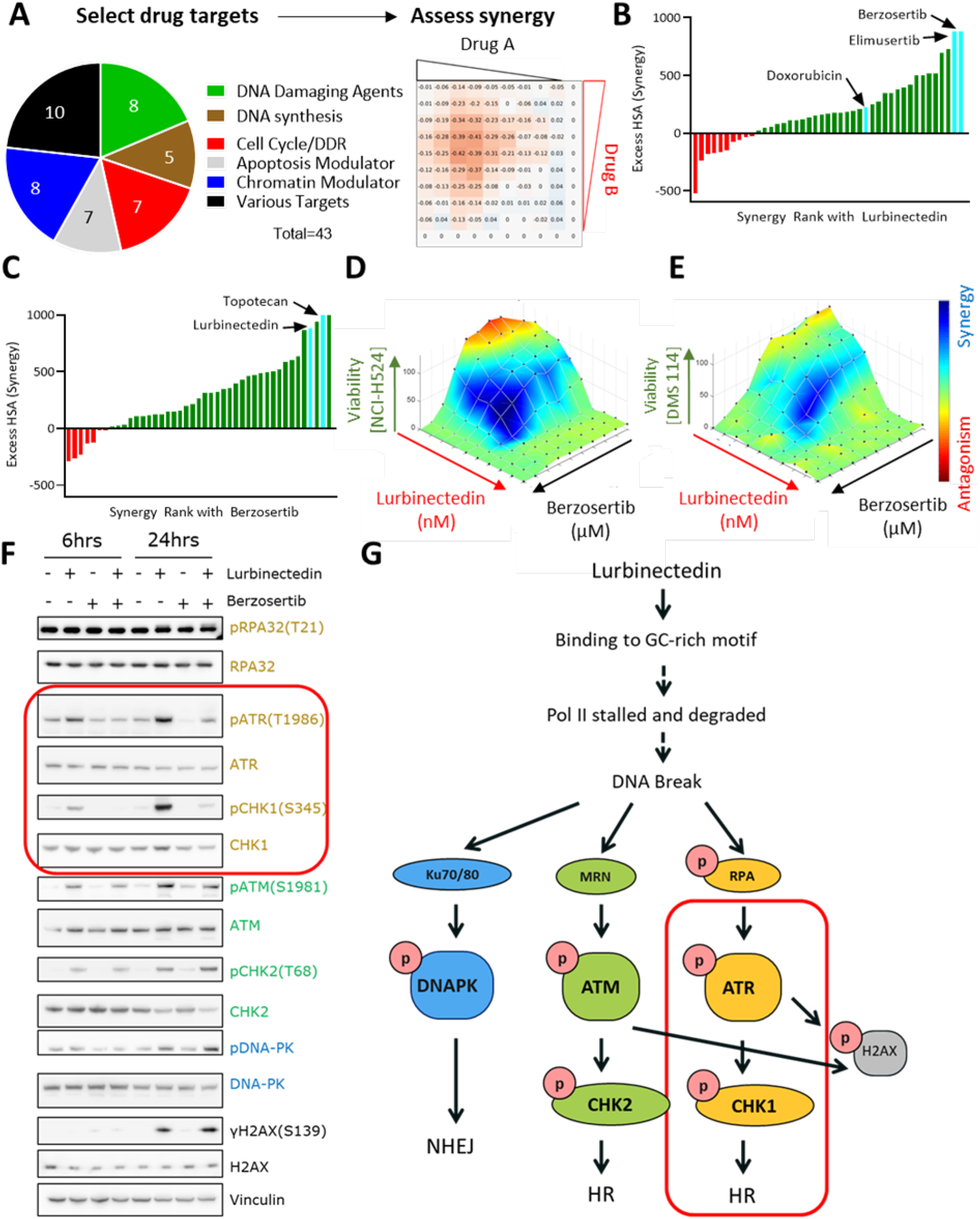
Lurbinectedin and berzosertib synergize in SCLC: **A)** A synergy screen was previously performed with 43 therapeutics targeting multiple pathways in combination with each other. NCI-H446 SCLC cells were treated with drugs in a 10x10 matrix format, viability was assessed using Cell Titer Glo and synergy was assessed using Highest Single Agent (HSA). **B)** Lurbinectedin synergy was ranked based on HSA synergy values. The highest synergy was observed with two ATR inhibitors, elimusertib and berzosertib, both of which displayed more synergy than doxorubicin. **C)** Berzosertib synergy with therapeutics ranked based on synergy. Topotecan and lurbinectedin both strongly synergized with berzosertib. **D**,**E)** Lurbinectedin and berzosertib synergized in SCLC cell lines DMS 114 and NCI-H524. Synergy of lurbinectedin and berzosertib was assessed by treating these drugs in a 10x10 matrix format for 72 hours in NCI-H524 (D) (HSA 410.7) and DMS 114 cells (E) (HSA 286.3). Synergy was calculated by adding HSA across all combination in the matrix (100 combinations, replicates=3, n=1). Synergy is denoted by blue and antagonism in red. **F)** Multiple DNA damage response pathways were activated by lurbinectedin. Treatment with the ATR inhibitor berzosertib specifically inhibited the activation of ATR and its downstream target CHK1 (both targets indicated by red box). DMS 114 cells were treated with lurbinectedin (1 nM) +/-berzosertib (1 µM) for 6 or 24 hours and targets were assessed by immunoblotting. **G)** Lurbinectedin activated all three key DNA damage response proteins, DNA-PK, ATR and ATM. Berzosertib is effective at inhibiting the activation of ATR and downstream ATR target CHK1 (indicated by red box), while not significantly impacting other pathways. berzosertib was also confirmed in SCLC cell lines NCI-H524 (HSA 410.7) and DMS 114 (HSA 286.3) (**Fig. 1 D, E**).

### Comet Assay

DMS 114 cells were plated at 500K cells per well in a 6 well plate, cells were treated +/-1nM lurbinectedin and +/-2µM berzosertib for 6 hours and collected. Comet assays was performed using the CometAssay Single Cell Gel Electrophoresis Assay (Trevigen, Gaithersburg, MD) according to manufacturer instructions. Images were captured using BioSpa Live Cell Analysis System (Biotek) and comet tail length was calculated using OpenComet (https://cometbio.org/), a plugin for the image processing program ImageJ.

### Immunoblotting

For immunoblotting experiments we utilized the following antibodies: -γH2AX(S139)-Cell Signaling #80312, H2AX-Sigma-Aldrich # 07627, pRPA32(T21)-Abcam #ab109394, RPA32-Cell Signaling #35869, - pATR(T1986)-Cell Signaling #58014 and #30632, ATR-Cell Signaling #13934, pCHK1(S345)-Cell Signaling #12302, CHK1-Cell Signaling #2360, pATM(S1981)-Abcam #ab81292, ATM-Cell Signaling #2873, pCHK2(T68)-Cell Signaling #2197, CHK2-Cell Signaling #6334, pDNA-PK(S2056)-Abcam# ab18192, DNA-PK-Abcam# ab32566, Vinculin-Sigma-Aldrich # V9131, Histone H3-Sigma-Aldrich #07-690.

### Immunofluorescence

Cells were fixed with 2% paraformaldehyde/PBS for 20 min at room temperature, washed 3 times PBS, cells were deposited on slide glass by cytospin, then put in pre-chilled 70% ethanol for 20 min, blocked with 5% BSA/PBSTT (PBS containing 0.5% Tween 20, 0.1% Triton X-100) for 30 min. Slides were incubated with primary antibodies for 2 hours and secondary antibodies for 1 hour at room temperature. Images were captured with a Zeiss LSM 780 confocal microscope.

### DNA Combing

DMS 114 cells were treated +/-1nM lurbinectedin and +/-2µM berzosertib for 6h followed by treatment with CIDU for 30 minutes and then IDU for 30 minutes and collected. DNA combing assay was performed as described previously [6, 22].

### Assessment of Mitotic Catastrophe by Fluorescence Microscopy

Using fluorescence microscopy, mitotic catastrophe was identified based on the characteristic morphologies of the DAPI-stained nuclei. Cells were seeded in 12-well plates on sterilized coverslips and treated the next day with either vehicle, 1nM lurbinectedin, 2µM berzosertib and combination of both for 6 hours. Nocodazole was added into media for 3 hours during the drug treatment (to enrich mitotic cells), then nocodazole and drugs were removed and fresh media was added for 45 min. After washing cells with cold PBS, fixation was done in 4% paraformaldehyde in PBS for 15 min at RT followed by permeabilization with 0.2% Triton-X 100/PBS for 15 min. After washing with PBS, cells were mounted with DAPI (VECTASHIELD, Vector Laboratories). Images of nuclei were captured with Zeiss LSM 880 confocal microscope with 63x objective lens.

### RNA analysis, NE calculation and enrichment analysis

RNA data and neuroendocrine score calculation for cell lines is publicly available on CellMiner [23]. Patient tumor and CDX datasets of SCLC were obtained from publicly available datasets (Integrated genomic and transcriptomic analysis of small cell lung cancer reveals inter-and intratumoral heterogeneity and a novel chemotherapy-refractory subtype, under revision, data available at dbGaP Study Accession: phs002541.v1.p1) [9, 24-26]. High MYC was determined in the patient and CDX datasets through assessing the expression of MYC, L-MYC and N-MYC, z-scoring them, and selecting the max z-scored expression, high-myc was defined as one stdev above mean. ssGSEA hallmark and neuroendocrine enrichment scores were computed using the GSVA R/Bioconductor package[13, 27, 28].

### Metaphase Spread

DMS 114 cells were plated at 1 million cells per 10cm plate, treated +/-1nM lurbinectedin and +/-2µM berzosertib for 6 hours and collected. Metaphase spread was performed as described https://ccr.cancer.gov/sites/default/files/metaphase_preparation_from_adherent_cells.pdf. Cells were imaged using a Leica Thunder Imager.

### Flow Cytometry

Cells were treated +/-1nM lurbinectedin and +/-2µM berzosertib for 6 hours, ethynyl deoxyuridine (EDU) at 100 µM was added for the last hour prior to collection collected. EdU was detected by flow cytometry (Click-iT EdU Alexa Fluor 647 Flow Cytometry Assay, Invitrogen), DNA using DAPI, and γH2AX was assessed by staining cells with JBW301-Millipore Sigma. Data were acquired using a BD LSRFortessa Flow Cytometer and analyzed using FlowJo.

## Results

### Drug screen identifies synergy of ATR inhibitors with lurbinectedin

To identify agents synergistically cytotoxic with lurbinectedin, we leveraged a previously reported drug-screen in the SCLC cell line NCI-H446 wherein lurbinectedin was combined with 43 FDA-approved drugs or agents in late stages of clinical development [6]. The compound library included DNA damaging agents and drugs that target mechanistically diverse pathways including DNA synthesis/metabolism, cell cycle or DNA damage repair, apoptosis, and chromatin remodeling (**Fig. 1A**). Synergy was assessed using the highest single agent (HSA) model [29], positive values denoting synergy and negative values antagonism. Lurbinectedin was most synergistic with DNA damaging agents, and drugs targeting cell cycle/DNA damage repair, and chromatin remodeling. In contrast, combinations with inhibitors of DNA synthesis and DNA metabolism were antagonistic, with the least synergy observed with the dihydrofolate reductase inhibitor pralatrexate (HSA -524.6) and the DNA polymerase inhibitor cytarabine (HSA -235.7) (**Table S1**).

Maximal cytotoxic synergy with lurbinectedin was observed for elimusertib (BAY1895344, HSA 881.2) and berzosertib (M6620, VX-970, VE-822, HSA 881.4), inhibitors of ATR, the main transducer of replication stress signaling and regulator of the cell cycle in response to DNA damage (**Fig. 1B, Table S1**). In comparison, both ATR inhibitors displayed greater synergy with lurbinectedin as compared to the topoisomerase II inhibitor doxorubicin (224.4 HSA, **Table S1**). Doxorubicin was previously reported to be synergistic with lurbinectedin in pre-clinical models, however this combination failed to improve survival compared with standard-of-care in patients with relapsed SCLC [30]. In a reciprocal screen, lurbinectedin was among the top four agents that showed maximal synergy with berzosertib (**Fig. 1C, Table S2**). The other top hits were inhibitors of key proteins involved in maintaining genomic stability including ataxia-telangiectasia mutated (ATM, AZD-0156)[31], WEE1 (MK-1775)[32] and topoisomerase I (TOP1) (topotecan) (**Table S2**). The combination of topotecan and berzosertib is being examined in clinical trials [5, 6] (NCT04768296, NCT03896503). Notably lurbinectedin was more potent than topotecan in 7/9 SCLC cell lines tested and thus may have improved efficacy (**Fig. S1A**). The synergy of lurbinectedin and

### The combination of lurbinectedin and berzosertib causes mitotic catastrophe

Lurbinectedin induced concentration and time-dependent activation of ATR, ATM and DNA-dependent protein kinase (DNA-PK), primary kinases that regulate DNA repair, and γH2AX, a marker of DNA DSBs [33] **(Fig. S1B)**. The addition of berzosertib reduced the activation of ATR and its downstream target CHK1, notably with little impact on ATM/CHK2 or DNA-PK (**Fig. 1F**). These results confirm that lurbinectedin-induced DNA damage activates multiple DNA damage sensing and repair pathways, but berzosertib specifically reduces the activation of the ATR-CHK1 axis **(Fig. 1G)**.

Given the critical role of ATR and its downstream target CHK1 in initiating cell cycle arrest in response to DNA damage, we investigated whether the synergy of lurbinectedin and berzosertib was cell cycle dependent. Lurbinectedin-induced DNA damage as measured by γH2AX occurred predominantly in the S-phase of cycling cells (**Fig. 2A, Fig. S1C**). Berzosertib alone or in combination with lurbinectedin reduced γH2AX in the majority of cells, but in a small fraction (∼5%) of the population caused a drastic increase in γH2AX. Therefore, when assessed using flow cytometry, co-treatment with berzosertib reduced lurbinectedin dependent S-phase γH2AX induction in the majority of cells leading to a decrease in median γH2AX signal in S-phase cells, but consistent with our immunoblotting (Fig 1F) caused an overall increase in average γH2AX signal in DMS 114 cells (**Fig. 2B, Fig. S1D**,**E**,**F**). We assessed γH2AX induction using immunostaining and immunoblotting in additional cell lines and found berzosertib overall inhibited lurbinectedin dependent γH2AX induction (**Fig. 2C, Fig. S2A**,**B**,**C**,**D**). Together, the immunoblotting, immunostaining and flow cytometry analysis indicated that berzosertib co-treatment reduces lurbinectedin-induced γH2AX. Reduced γH2AX signal in the majority of cells was not due to a decrease in DNA damage as the combination of lurbinectedin and berzosertib induced more DNA breaks as assessed by alkaline comet assay than either agent alone (**Fig. 2D,E**). Treatment with lurbinectedin reduced DNA replication as indicated by a decrease in 5-ethynyl-2”-deoxyuridine (EdU) incorporation, an effect which was largely rescued by berzosertib, indicating that the decrease in DNA replication was ATR dependent (**Fig. 2F, Fig. S1G**).

**Fig. 2.**
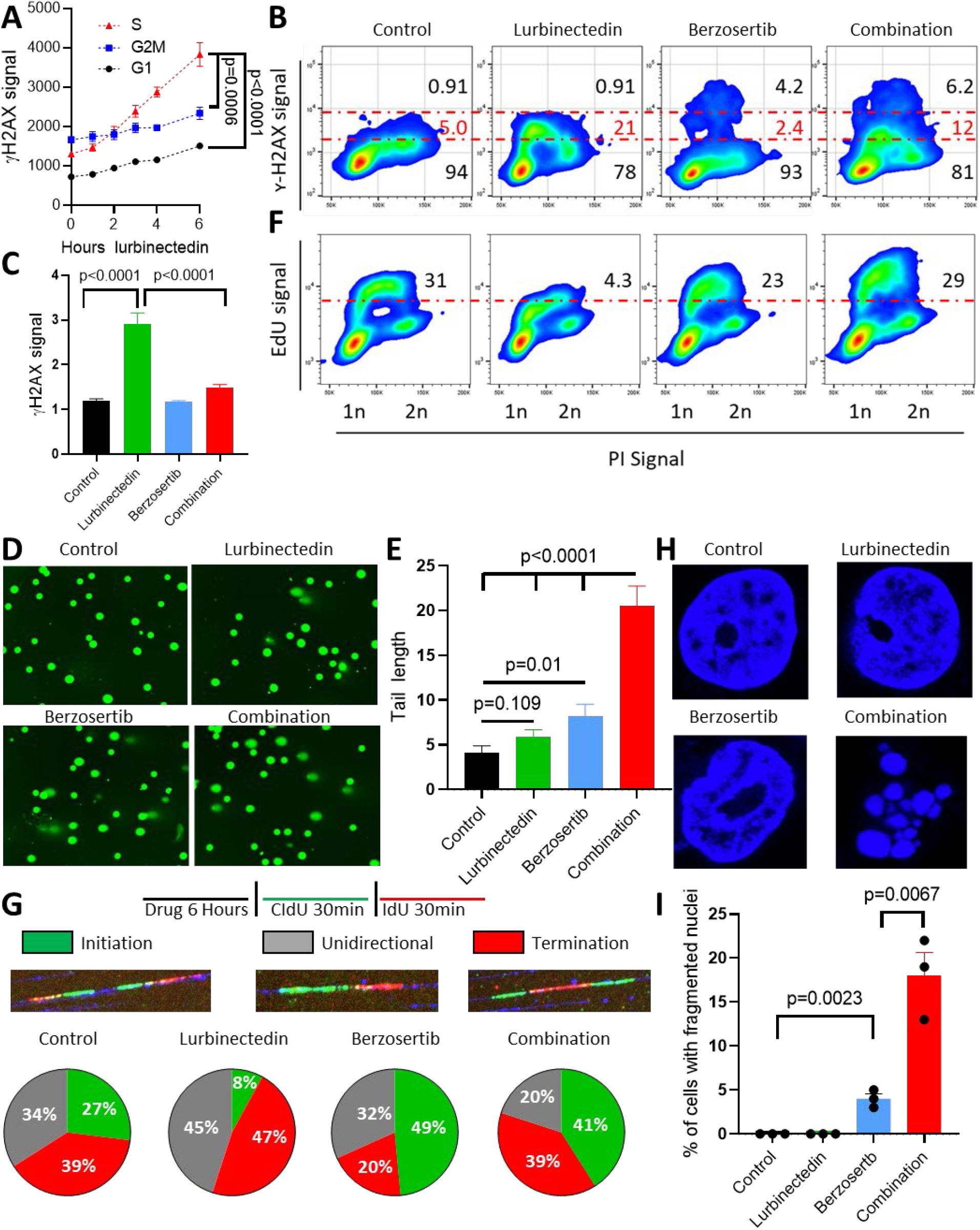
Berzosertib causes continued cell cycle progression and induces mitotic catastrophe in the presence of lurbinectedin: **A)** Lurbinectedin induced changes to γH2AX across different phases of the cell cycle, with the greatest degree of activation in S phase. DMS 114 cells were treated with 1 nM lurbinectedin for 1 to 6 hours and γH2AX induction was assessed using flow-cytometry, four replicates of 10,000 cells for each timepoint were assessed, error bars represent SEM, n=3. Cell cycle was assessed using propidium iodide. **B)** Lurbinectedin treatment increased γH2AX induction, while berzosertib treatment caused a small portion of cells to display increased γH2AX signal. Berzosertib co-treatment with lurbinectedin caused a decrease in γH2AX expression for the majority of cells leading to a decreased median expression of γH2AX, however an increase in γH2AX in a smaller portion of cells led to an increased average expression (averages and medians are shown in Fig. S1D,E,F). DMS 114 cells were treated +/-1 nM lurbinectedin +/-2 µM berzosertib for 6 hours γH2AX induction was assessed using flow-cytometry. 10mM EdU was added for the last hour prior to collection (assessed Fig 2F). Numbers on the graph indicate the average percent of cells across 4 replicates of 10,000 cells with low, medium or high γH2AX n=3. **C)** Lurbinectedin treatment increased γH2AX induction which was reduced with berzosertib treatment. γH2AX induction was assessed using immunofluorescence in DMS 114 cells treated +/-1 nM lurbinectedin +/-2µM berzosertib for 6 hours. Quantification is from 100-150 cells per treatment, with error bars indicating SEM, comparisons were made using an unpaired two-tailed student”s t test in PRISM, n=3. Representative images for this experiment are in Fig. S2A. **D/E)** Combination of lurbinectedin and berzosertib caused DNA damage. DMS 114 cells were treated +/-1 nM lurbinectedin +/-2 µM berzosertib for 6 hours and total DNA damage was assessed using an alkaline comet assay. These data represent average tail length for each group with 150-250 cells measured per group, n=3 error bars represent SEM, p values represent unpaired two-tailed student”s t tests performed in PRISM. **F)** Lurbinectedin treatment caused a decrease in DNA replication as assessed by EdU incorporation while berzosertib co-treatment largely rescued this phenotype. Numbers represent the average percent of cells across 4 replicates with high EdU incorporation indicates functional DNA replication, n=3. These cells are the same as assessed in Fig 2B. **G)** Lurbinectedin treatment inhibited fork initiation, an effect which was rescued by the addition of berzosertib. DMS 114 cells were treated +/-1 nM lurbinectedin +/-2 µM berzosertib for 6 hours and using DNA combing we assessed motifs of incorporation of IdU and CIdU indicating initiation, termination or unidirectional DNA forks. Forks were assessed and verified individually to avoid algorithmic assessment, with 100-200 forks quantified per group n=2. **H**,**I)** Berzosertib treatment alone caused a small portion of cells to undergo mitotic catastrophe, however cotreatment of lurbinectedin and berzosertib caused a large portion of cells to undergo mitotic catastrophe. DMS 114 cells were treated +/-1 nM lurbinectedin +/-2 µM berzosertib for 6 hours with the final three hours being in the presence of nocodazole followed by release from all drugs and 45 minutes of continued growth. Cells which had undergone mitotic catastrophe were assessed as those which were multinucleated. Representative images (H) and quantification of percent of cells which underwent mitotic catastrophe (I). Graph represents average of three separate experiments with statistics determined from assessing 100 nuclei per group, error bars represent SEM, p values are indicative of unpaired two-tailed student”s t tests performed in PRISM.

Next, we used molecular combing (DNA fiber assays) to visualize single replicons and measure the impact of lurbinectedin and ATR inhibition on replication dynamics. Control DMS 114 cells displayed roughly equal distribution of initiation (27%), termination (39%) and unidirectional (34%) forks. Lurbinectedin treatment suppressed replication fork initiation by 3-fold while slightly increasing termination and unidirectional forks (1.2-fold and 1.3-fold respectively). These results were in agreement with our previous data indicating lurbinectedin treatment inhibited DNA synthesis. Berzosertib treatment increased initiation forks by 1.8-fold compared with control, consistent with previous work demonstrating that ATR inhibition causes the unscheduled firing of dormant origins [4], with a minimal impact on termination and unidirectional forks. In combination, berzosertib treatment rescued the suppression of replication fork initiation caused by lurbinectedin, with a 5-fold increase in initiation forks as compared to lurbinectedin treatment, also reducing termination and unidirectional forks to a lesser extent (0.82-fold and .45-fold respectively) **(Fig. 2G)** [34]. These results demonstrated that berzosertib leads to unscheduled replication origin firing and continued DNA replication even in the presence of lurbinectedin-induced DNA damage.

In addition to its crucial role in DNA damage and replication stress response, ATR also promotes accurate chromosome segregation during mitosis [33]. We assessed metaphase spreads to determine drug treatment-induced changes to chromosomal integrity during mitosis. Berzosertib monotherapy led to abnormal metaphase spreads, with individual chromosomes failing to segregate appropriately (**Fig. S2E**,**F**). This phenotype was recapitulated on treatment with barasertib which inhibits Aurora Kinase B, a downstream target of ATR critical for accurate chromosomal segregation (**Fig. S2G**). Lurbinectedin alone had little effect on chromosomal integrity at metaphase. Lurbinectedin-berzosertib combination treatment did not impact the frequency of segregation defects caused by berzosertib treatment, but the combination significantly increased the percentage of mitotic cells with multiple breaks in chromosomes (**Fig. S2E**). We assessed post-mitotic viability in order to determine if the chromosome breaks induced by co-treatment effected mitotic competence. Both lurbinectedin and berzosertib had little effect on post-mitotic viability by themselves. The combination however, significantly increased post-mitotic death as indicated by multi-nucleation (**Fig. 2H,I**). Together, these results demonstrate that berzosertib augments DNA damage caused by lurbinectedin and allows cells to progress to mitosis with unrepaired DNA damage ultimately leading to mitotic catastrophe and cell death.

### *ERCC5*/XPG and SLFN11 as critical biomarkers of response to lurbinectedin

Similar to the structurally related trabectedin, adducts generated by binding of lurbinectedin to the DNA minor groove are recognized by the nucleotide excision repair (NER) pathway. The NER complex then becomes trapped on the DNA, ultimately leading to the formation of irreversible single-strand-breaks, and consequently cell death [19, 35]. Homologous recombination (HR) is critical for the repair of lurbinectedin induced DNA damage, and loss of the HR pathway increases sensitivity to lurbinectedin [35]. Isogeneic models using the chicken B cell line DT40 cell have previously been utilized to demonstrate the importance of NER and HR pathways for lurbinectedin efficacy [35]. We examined the contribution of these DNA repair mechanisms to the efficacy of lurbinectedin, berzosertib, and the combination utilizing isogenic DT40 models. Consistent with previous findings, DT40 cells with knockout (KO) of the NER pathway mediator *ERCC5*/XPG were approximately 10-fold more resistant to lurbinectedin. Alternatively, *BRCA2*-KO DT40 cells were approximately 6-fold more sensitive to lurbinectedin (**Fig. S3A**). Berzosertib efficacy was not significantly altered in the knockouts (**Fig. S3B**).

ATR signaling and ATR induced cell cycle arrest are critical for efficient HR repair [33]. To confirm that berzosertib was inhibiting HR competency we utilized U2OS cells with a stably integrated GFP HR reporter [36]. We confirmed that berzosertib inhibited HR competency at 7-fold lower concentrations than that required for inhibiting viability (**Fig. S3C**,**D**). If lurbinectedin-berzosertib synergy was due to reduced HR competency from berzosertib, synergy would be decreased in the *BRCA2*-KO. Supporting this hypothesis, *BRCA2*-KO cells displayed slightly decreased synergy. Consistent with the importance of the NER pathway for lurbinectedin efficacy, synergy was even more markedly reduced with *XPG*-KO cells (**Fig. S3E**). To standardize the assessment of overall combination efficacy, we defined lurbinectedin IC_50_ in the presence of 500nM of berzosertib (a concentration where lurbinectedin IC_50_s are reduced in all synergistic models tested and HR is inhibited) as representative of overall combination efficacy.

Lurbinectedin-berzosertib combination displayed similar combination efficacy in the *BRCA*-2 KO cell line as compared to control. However, the *XPG*-KO cell line maintained resistance even in the presence of 500nM berzosertib leading to reduced combination efficacy (**Fig. S3F**). These results show that lurbinectedin-berzosertib synergy is only partially mediated by berzosertib inhibiting HR competency, and that the efficacy of lurbinectedin alone and combination efficacy are both dependent on NER competency and *ERCC5*/XPG expression.

A recent report has indicated that cells with high expression of SLFN11, another important mediator of DNA damage response, are significantly more sensitive to lurbinectedin [37]. SLFN11 destabilizes paused replication forks following DNA damage causing cell death. Consequently, loss of SLFN11 leads to broad resistance to a variety of DNA damaging therapeutics [38]. We found that SLFN11-KO DMS 114 cells were approximately 4-fold more resistant to lurbinectedin than parental cells, while there was little difference in sensitivity to berzosertib **(Fig. S3G**,**H**). Consistent with previous findings demonstrating the ability of ATR inhibition to re-sensitize SLFN11-KO cells to DNA damaging agents [38], synergy was increased in SLFN11-KO cells, and SLFN11-KO cells displayed similar combination efficacy as the control cells **(Fig. S3I**,**J**). As berzosertib rescued lurbinectedin efficacy in SLFN11-KO cells and not in XPG-KO cells we conclude that *ERCC5*/XPG is mechanistically required for lurbinectedin activity even in the presence of an ATR inhibitor, whereas SLFN11 is not. These results are in agreement with SLFN11 inducing lethal replication fork instability in response to DNA damaging agents independently of ATR and acting in parallel with the ATR-mediated S-phase checkpoint [39].

Next, we assessed the efficacy of lurbinectedin, berzosertib and the combination of both agents in a 10x10 matrix format (100 conditions) across a panel of nine SCLC cell lines spanning the spectrum of neuroendocrine differentiation and lineage transcription factors (**Table 1**) [16]. Lurbinectedin was highly potent across SCLC cell lines with IC_50_s ranging from 0.01–0.38 nM, while berzosertib IC_50_s ranged from 0.33-5.03 µM (**Table 1**). Consistent with findings in isogenic models described above, we observed trends toward increased sensitivity to lurbinectedin in cells with high *SLFN11* or *ERCC5*/XPG expression (**Fig. S4A**). We assessed the IC_50_ of lurbinectedin in the presence of 500 nM berzosertib, a concentration which effectively inhibits HR, and has little effect on viability by itself in most models. Increased *ERCC5*/XPG expression significantly correlated with improved combination efficacy, while higher *SLFN11* expression trended towards increased combination efficacy (**Fig. 3A, Table 1, Table 2, Fig. S4A**). Overall, studies in isogenic systems and human SCLC cell lines demonstrate that SLFN11, HR and *ERCC5*/XPG are determinants for the response to lurbinectedin monotherapy, with *ERCC5*/XPG being a critical determinant of response to the lurbinectedin-berzosertib combination i.e. combination efficacy (**Table 2 Fig. S4B**).

**Fig. 3.**
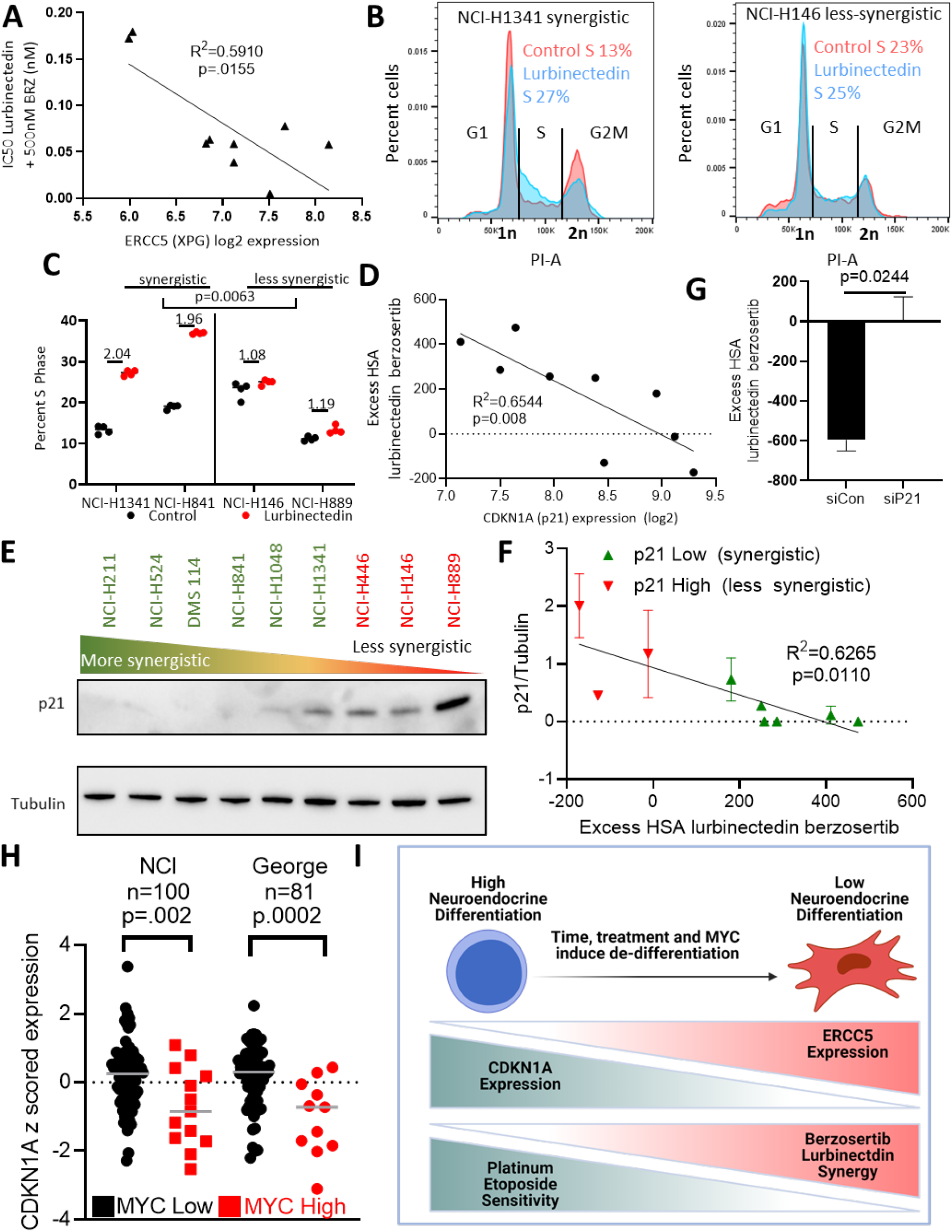
p21 inhibits lurbinectedin and berzosertib synergy, with greater synergy expected in recalcitrant MYC driven non-NE cells: **A)** *ERCC5* (XPG) expression inversely corelated with combination efficacy, i.e. when the expression of *ERCC5* (XPG) was high the combination was more effective. The IC_50_ of lurbinectedin in the presence of 500 nM berzosertib (combination efficacy) was determined in 9 cell lines after assessing all 9 lines using a 10x10 matrix of lurbinectedin and berzosertib combinations (replicates =3, n=1), Pearson correlation was assessed in PRISM. **B**,**C)** Lurbinectedin increased S phase arrest as assessed by propidium iodide staining. Synergistic cell lines had a greater increase in S phase than less synergistic lines. Synergistic (NCI-H841 and NCI-H1341) and less synergistic (NCI-H146 and NCI-H889) cell lines were treated +/-1nM lurbinectedin +/-2µM berzosertib. Quantification in C is representative of 4 replicates of 10,000 cells per group n=3, with fold change in S phase cells being compared between synergistic and less synergistic cell lines using an unpaired two-tailed student”s t test in PRISM. These data are also described in Fig. S4C. **D)** *CDKN1A* (p21) RNA expression inversely correlated with synergy of lurbinectedin and berzosertib, indicating that *CDKN1A* (p21) could potentially inhibit synergy. Pearson correlation was assessed in PRISM. **E**,**F)** Synergy was lower in cells which displayed higher expression of p21 protein. (E) Cell lines were ordered by HSA synergy score (most to least synergy left to right) and p21 protein expression was assessed by immunoblotting. (F) Quantitation of p21 protein compared to control Tubulin across two independent experiments with fresh samples plotted against synergy values, error bars represent SD, Pearson correlation was assessed in PRISM. **G)** Knockdown of CDKN1A (p21) in NCI-H889 cells led to increased synergy. NCI-H889 cells were treated with siRNA against control or *CDKN1A* (p21), followed by dosing in a 10x10 matrix format in triplicate with lurbinectedin and berzosertib and collection after 72 hours. HSA synergy was determined and summed across the 10x10 matrix, graph represents the average of 2 independent experiments of 10x10 matrixes in triplicate, error bars =SD, unpaired two-tailed student”s t test performed in PRISM. **H)** High MYC family member patient samples had lower CDKN1A (p21) expression consistent with high MYC family member expression causing decrease CDKN1A (p21). In two independent SCLC datasets, *MYC, MYCL* and *MYCN* expression were z scored (within each database) and the max MYC family member z score expression was determined for each sample. Those samples which were greater than one standard deviation above average were considered to be high MYC family member expressing. P values are indicative of unpaired two-tailed student”s t test assessed in PRISM. **I)** As SCLC progresses cancer cells progress from a NE differentiated state to a non-NE state. These non-NE cells have a lower expression of *CDKN1A* (p21) and a higher expression of *ERCC5* (XPG). These are characteristics which make them less responsive to the standard first line platinum/etoposide regimen, however this should make them more sensitive to the combination of lurbinectedin and berzosertib.

### The G1/S checkpoint is a critical determinant of lurbinectedin-berzosertib synergy, consequently *CDKN1A*/p21 is a biomarker of reduced synergy

The lurbinectedin-berzosertib combination was synergistically cytotoxic in six of nine SCLC cell lines assessed (HSA 180.5 to 474.0) and additive (HSA -11.9 where 0 is additive) or antagonistic (HSA -128.3 to -171.3) in the others (**Table 1**). Synergy i.e., HSA or the difference between lurbinectedin IC_50_s +/-berzosertib, did not correlate with sensitivity to either drug or to the expression of *SLFN11* or *ERCC5*/XPG (**Fig. S4A**). As lurbinectedin and berzosertib greatly effected DNA replication and mitotic division, we assessed whether synergy was dependent on cell cycle dynamics. Cell lines with higher synergy (NCI-H841 and NCI-H1341) upon exposure to lurbinectedin had significantly greater S-phase arrest which was ameliorated with co-treatment with berzosertib as compared to less synergistic cell lines (NCI-H146 and NCI-H889) (**Fig. 3B,C**) (**Fig. S4C**). Therefore, we hypothesized that cells deficient in the G1/S checkpoint could be uniquely sensitive to the combination of lurbinectedin-berzosertib.

SCLC is characterized by loss of RB1 [9], the predominant regulator of G1/S transition leading to increased reliance on cyclin dependent kinase inhibitor p21 (*CDKN1A*) for control of the G1/S checkpoint [40]. High expression of *CDKN1A* RNA was associated with reduced lurbinectedin-berzosertib synergy (**Fig 3D, Table 1, Table 2**). Furthermore, p21 protein expression, which was highly correlated with *CDKN1A* RNA expression (**Fig. S4D**), was also associated with decreased synergy (**Fig. 3E,F**). Supporting our hypothesis that p21-mediated G1 arrest could lead to reduced sensitivity to lurbinectedin-berzosertib combination, thymidine enforced G1 cell cycle arrest led to a significant reduction in efficacy of the combination (**Fig. S4E**). Small interfering RNA (siRNA) knockdown of p21 in the least synergistic cell line NCI-H889 resulted in a significant increase in synergy (**Fig. 3G**). These data are consistent with previous work in which p21 levels predicted reduced sensitivity to agents targeting downstream targets of ATR, CHK1 and Wee1 [40].

ATM-dependent activation of p53 followed by p53-dependent p21 up-regulation is a canonical pathway that regulates p21 expression in response to DNA damage [41]. *TP53*/p53 is mutated in most SCLC tumors [9], suggesting frequent abrogation of the ATM-p53-p21 axis in SCLC. To determine the impact of ATM-p53-p21 pathway on lurbinectedin-berzosertib synergy, we utilized two cell lines which exhibited low synergy and high expression of *CDKN1A* which were *TP53* mutated (NCI-H889), or *TP53* intact (NCI-H146). Upon lurbinectedin treatment, p21 protein levels were reduced in NCI-H889 cells, whereas NCI-H146 cells displayed an increase in p21 (**Fig. S5A**). P21 decrease in NCI-H889 cells is expected as lurbinectedin inhibits RNA Pol-II and, due to the rapid turnover rate of p21 protein and RNA, inhibition of RNA Pol-II causes rapid decreases in p21 [42]. Increased p21 in NCI-H146 cells is consistent with the ability of both DNA damaging agents and transcription inhibitors to increase p21 expression in an ATM-p53-p21 axis dependent manner [41]. Consistent with the importance of ATM in the *TP53* intact setting, addition of the ATM inhibitor KU60019 [43] significantly increased the synergy of lurbinectedin-berzosertib in H146 cells. Conversely in NCI-H889 cells ATM inhibition was less effective at increasing synergy (**Fig. S5B**,**C**).

Together, our *in-vitro* data show that high *CDKN1A*/P21 expression irrespective of *TP53* status predicts decreased synergy of lurbinectedin-berzosertib i.e., less impact on lurbinectedin IC_50_s with the addition of berzosertib, while higher *ERCC5*/XPG expression predicts greater overall sensitivity i.e., decreased lurbinectedin IC_50_s in the presence of berzosertib. We confirmed these results in two organoid models of SCLC (**Fig. S5D**,**E**,**F**,**G**). We propose *CDKN1A*/P21 as the primary negative determinant of synergy for lurbinectedin-berzosertib while *ERCC5*/XPG expression is required for lurbinectedin induced DNA damage and determines combination efficacy (**Table 2**).

### MYC expression and non-NE differentiation are associated with lurbinectedin-berzosertib synergy

SCLC is characterized by NE differentiation which decreases as the tumors progress and following chemotherapy [14, 44]. NE differentiation as characterized by a previously validated 50-gene signature [13] negatively correlated with lurbinectedin-berzosertib synergy, with the highest synergy observed in non-NE SCLC cells (**Fig. S5H**). MYC, as a driver of both non-NE differentiation and decreased *CDKN1A* expression [44, 45] could potentially cause the increased synergy of lurbinectedin-berzosertib in non-NE cells. In metastatic small cell patient tumors (n=100), *MYC* expression was negatively correlated with NE differentiation (**Fig. S6A**). In additional primary SCLC patient tumor, circulating tumor cell derived xenograft, and patient derived xenograft (PDX) datasets, we observed that samples overexpressing *MYC* paralogues (*MYC, MYCL* or *MYCN*) consistently displayed significantly lower *CDKN1A* expression (**Fig. 3H, Fig. S6B**). Similar results were observed in SCLC cell lines where *MYC* expression negatively correlated with *CDKN1A* expression and NE differentiation (**Fig. S6 C**,**D**).

Confirming the recalcitrance of high MYC non-NE SCLC tumors [46], out of 50 hallmark genesets we found that the two genesets predictive of MYC activity, MYC-targets-V1 and MYC-targets-V2, were respectively the first and third most highly associated with platinum resistance in SCLC patients [28] [47] (**Fig. S6E**). MYC-targets-V1 significantly differentiated platinum/etoposide sensitivity (**Fig. S6F**,**G**), with MYC-low patients displaying hazard ratios of 0.2130 and 0.1830 at the median and quartile levels respectively for platinum resistance (response duration of <90 days to carboplatin etoposide therapy). These data indicate that tumors which were MYC-low were ∼5 fold less likely to be resistant to carboplatin/etoposide therapy as compared to MYC-high tumors. Interestingly, SCLC cell lines with enrichment of MYC-targets-V2 were associated with increased lurbinectedin-berzosertib combination efficacy (**Fig. S6H**). NE differentiation negatively correlated with *ERCC5*/XPG expression in multiple clinical datasets, with non-NE tumors displaying higher *ERCC5*/XPG expression (**Fig. S6I**,**J**). Together these data indicate that MYC driven, non-NE tumors are resistant to platinum-based chemotherapy, but they may be uniquely sensitive to lurbinectedin-berzosertib due to high *ERCC5*/XPG and low *CDKN1A*/p21 expression (**Fig. 3I, Table2**).

### In-vivo synergy and schedule dependence of lurbinectedin-berzosertib combination

In our previous work and current clinical trials testing the tolerability and efficacy of topotecan and berzosertib we have observed efficacy using a dosing regimen in which topotecan is treated daily for five days while berzosertib is dosed on days two and five of seven-day cycles [6]. Adjusting this model for the standard dosing of lurbinectedin we assessed the efficacy of lurbinectedin and berzosertib in mouse models of SCLC dosing lurbinectedin on day one, followed by berzosertib on days two and five of seven-day cycles. In a patient derived xenograft (PDX) model of SCLC, PDX-06, lurbinectedin was extremely effective by itself almost completely inhibiting all tumor growth. Likely due to the high efficacy of lurbinectedin alone, we observed minimal increased activity with the addition of berzosertib (**Fig. 4A**). We assessed a separate cohort mice treated in the same manner for target engagement 24 hours after dosing and found that lurbinectedin appeared to cause an increased in p-CHK1, a downstream target of ATR, while berzosertib co-treatment reduced p-CHK1 activation (**Fig. 4B,C**, **Fig. S7A**). Even though in initial testing mice treated for a single cycle displayed no toxicity, the combination was toxic with repeated dosing. In particular, significant damage at tail veins of lurbinectedin and combination dosed mice was observed; and the majority of these mice were sacrificed due to weight loss (**Fig. S6K**).

**Fig. 4.**
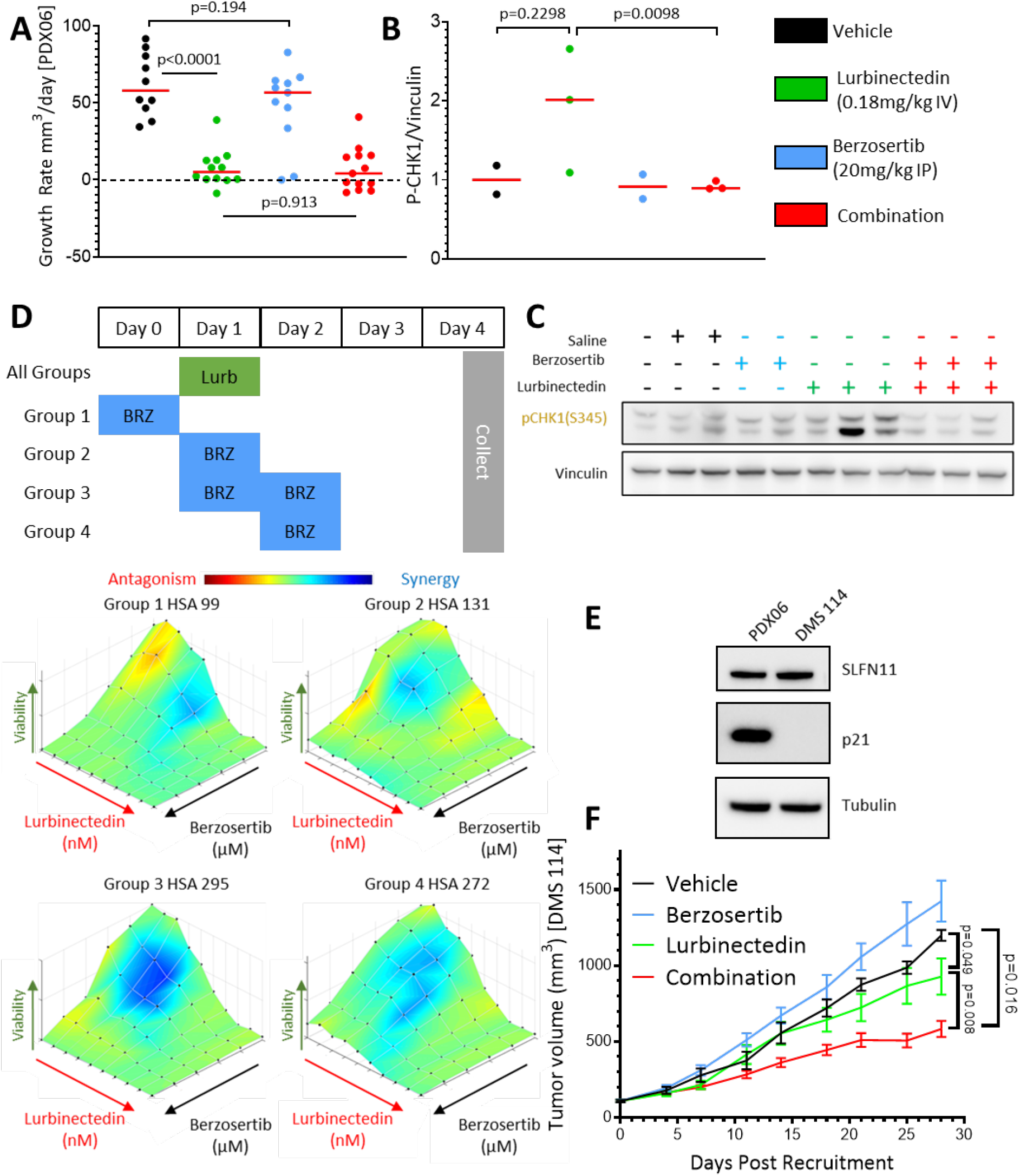
Berzosertib improves lurbinectedin efficacy *in-vivo*.: **A)** Lurbinectedin was very efficacious and the addition of berzosertib was did not significantly improve the almost complete inhibition of tumor growth caused by lurbinectedin in a PDX mouse model of SCLC (PDX-06), n=10-13 mice per group, red bar indicates median, p values are indicative of unpaired two-tailed student”s t test calculated in PRISM. Tumor-bearing mice were treated with lurbinectedin (0.18 mg/kg IV) and berzosertib (20mg/kg IP) in a format mirroring our clinical trial with topotecan/berzosertib in SCLC, lurbinectedin day 1, berzosertib days 2 and 5 of a 7-day cycle. Tumor growth rates were calculated as the difference between the initial tumor size and the final tumor size after death of the mouse due to toxicity or progression of tumor divided by the total number of days treated. **B**,**C)** We determined that lurbinectedin trended towards increasing p-chk1 while berzosertib co-treatment significantly reduced p-chk1 activation *in-vivo* indicating target engagement. Tumors from mice with the same tumor type (PDX-06) as in 4A which were collected 24 hours after being dosed with the indicated drugs. P values are indicative of unpaired two-tailed student”s t test calculated in PRISM. **D)** Lurbinectedin synergy was maximal in DMS 114 cells when treated day 1 with lurbinectedin and days 1 and 2 with berzosertib. DMS 114 cells were treated with lurbinectedin and berzosertib in a 10x6 matrix format replicates =4, n=1. All groups were treated for 24 hours with lurbinectedin, while group 1 was pretreated with berzosertib, group 2 was co-treated with berzosertib, group 3 was co and post treated with berzosertib, and group four was post treated with berzosertib for 24 hours. At the end of 5 days cells were collected and synergy was assessed across the matrixes. **E)** PDX-06 (less synergistic) and DMS 114 (more synergistic) cells had equivalent SLFN11 while DMS 114 had less p21. p21 and SLFN11 protein expression was assessed using immunoblotting. **F)** Berzosertib co-treatment improved lurbinectedin efficacy in a DMS 114 xenograft mouse model of SCLC. Lurbinectedin was dosed at (0.18mg/kg intravenous) and berzosertib (50mg/kg oral) for four 7 day cycles in dosing regiments consistent with 4D n=10 mice per group. Consistent with our results in 4D the greatest degree of increased and overall efficacy was seen in group 3 (lurbinectedin d1, berzosertib d1,2), p values are indicative of paired two-tailed student”s t test calculated in PRISM. The other groups are in displayed in Fig.S7 H,I.

To better assess combination efficacy, we utilized a more aggressive PDX model PDX-03. PDX-03 displayed reduced *CDKN1A*/p21 and *SLFN11* expression and increased aggressiveness as compared to PDX-06, and thus we expected it to be more resistant to lurbinectedin and to display increased synergy for the combination (**Fig. S6L**). To reduce toxicity, we doubled the volume of lurbinectedin dosed for tail vein injections (100µl to 200µl). We found in this model that lurbinectedin and combination efficacy was significantly reduced, potentially due to increased overall tumor aggressiveness in the PDX-03 model **(Fig. S7B**,**C, Fig. S6L)**. Although toxicity was reduced in the PDX-03 model, several mice in lurbinectedin and combination treatment arms still succumbed to toxicity as opposed to tumor burden (**Fig. S7D**). Due to the reduced efficacy and toxicity we were able to collect tumors from control and treatment arms and stain them for markers of proliferation and apoptosis. We found that while the combination had little increased efficacy as compared to lurbinectedin alone, it trended towards decreased ki-67 staining and increased cleaved caspase 3 and necrosis in tumor samples as compared to either lurbinectedin or berzosertib alone (**Fig. S7E**). We determined that the toxicity of the single agent and combination may potentially be due to the strain of mice, as severe combined immunodeficient (SCID) are inherently more sensitive to DNA damaging agents[48], and thus decided to utilize a nude mouse model which would likely have reduced toxicity.

Due to the limited *in-vivo* efficacy, we decided to re-assess the dosing strategy utilized. We assessed the synergy of lurbinectedin and berzosertib with berzosertib dosed either prior to lurbinectedin, co-treated with lurbinectedin, post-treated after lurbinectedin, or co-treated with and post-treated after lurbinectedin (lurbinectedin day 1, berzosertib day(s) 0/1/2/1+2). We found in two cell lines (NCI-H446 and DMS 114) that the greatest degree of synergy with lurbinectedin and berzosertib occurred when berzosertib was co-treated with lurbinectedin and then maintained after the removal of lurbinectedin (**Fig 4D, Fig. S7F**). As DMS 114 cells had a lower NE score than NCI-H446 cells we chose the DMS 114 cells to represent the recalcitrant non-NE subtype for our mouse model. We were able to derive a cell line from the NE PDX mouse model PDX-06, although unfortunately PDX-03 cells did not take to culture as well. We determined that PDX-06 cells expressed similar levels of SLFN11, but significantly higher p21 than the non-NE DMS 114 cells (**Fig. 4E**). Based on these features we expected the combination to be more synergistic in DMS 114 cells as opposed to PDX-06 cells. Indeed, compared with PDX-06 cells, DMS 114 cells were more resistant to lurbinectedin alone, but displayed significantly higher synergy for the lurbinectedin-berzosertib combination (**Fig. S7F**,**G**). In a xenograft model of DMS 114 cells in nude mice, the addition of berzosertib to lurbinectedin significantly increased efficacy (**Fig. 4F**). Consistent with our cell based work the greatest degree of synergy and activity was observed with lurbinectedin treatment on day 1 and berzosertib dosed on days 1 and 2; this combination was well tolerated (**Fig. 4F, Fig. S7H**,**I**). Of note, post-treatment with berzosertib was the least synergistic in our mouse model (**Figure S7H**) and sub-optimal in both cell lines (**Fig 4D, Fig. S7F)**, potentially explaining the lower degree of synergy in our PDX models in which berzosertib was post-treated. Notably, plasma drug levels achieved in the xenograft experiments were comparable to those of human patients (**Fig. S7J, K**) [6, 49].

Overall, lurbinectedin causes DNA damage in an *ERCC5*/XPG dependent manner, at which point cells can either be arrested in a *CDKN1A*/p21 (G1-phase checkpoint) or ATR (S-phase checkpoint) dependent manner. Cells arrested in S-phase due to ATR activation will either die via SLFN11-dependent lethal replication fork instability or resolve DNA damage through HR and continue through the cell cycle. The addition of berzosertib causes cells that would normally halt cell cycle progression in S-phase to continue through the cell cycle despite DNA damage and undergo mitotic catastrophe. XPG-dependent induction of DNA damage or p21 dependent G1-arrest occur upstream of ATR-activation and thus are unaffected by berzosertib treatment. As such *ERCC5*/XPG and *CDKN1A*/p21 are both critical determinants of lurbinectedin-berzosertib efficacy (**Fig. 5, Table 1, Table 2**).

**Fig. 5.**
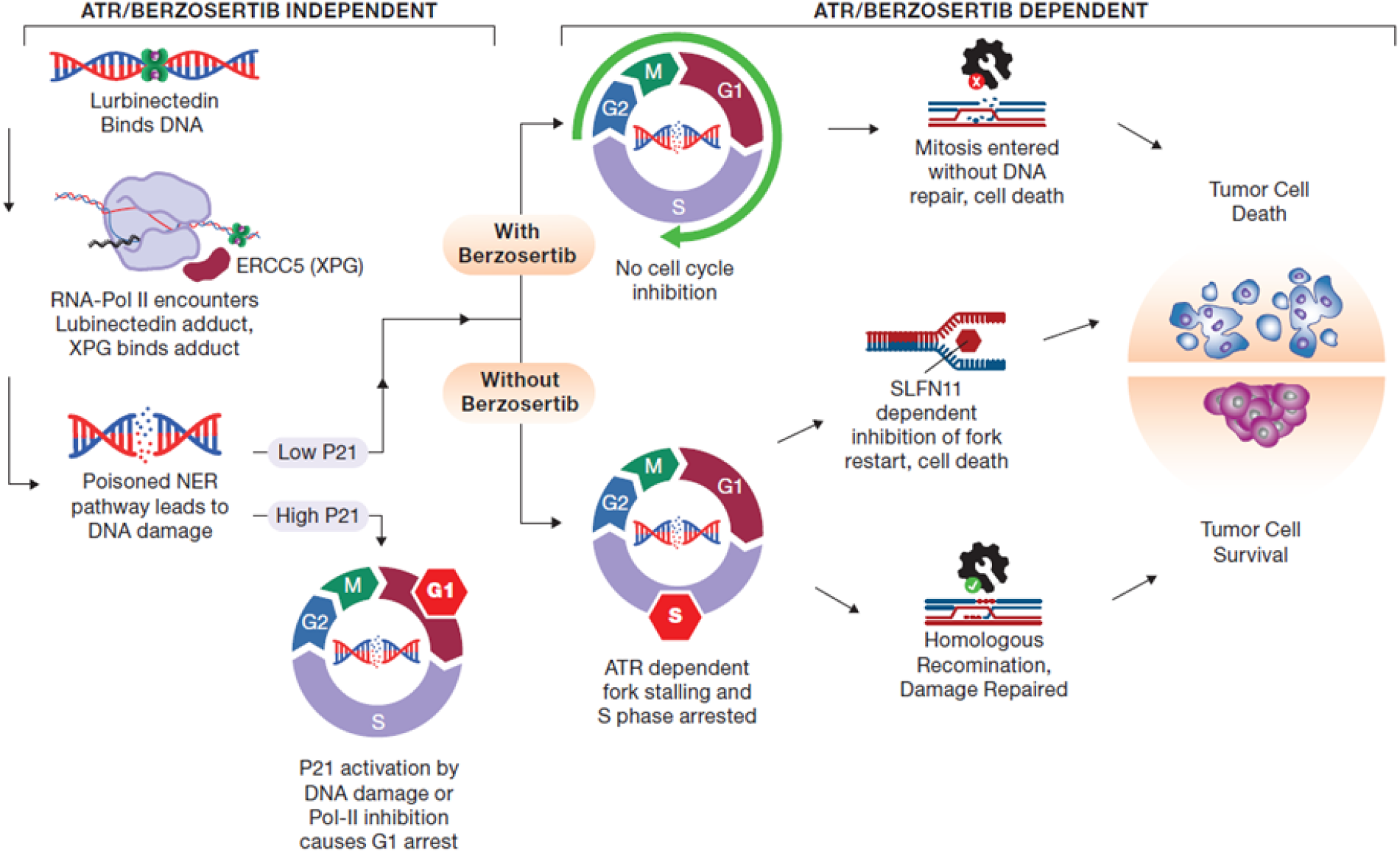
Berzosertib synergizes with lurbinectedin through inhibiting S phase arrest and inducing mitotic catastrophe: Lurbinectedin binds to DNA and then induces DNA damage through transcription coupled nucleotide excision repair poisoning in an XPG (ERCC5) dependent manner. p21 (CDKN1A) activation can lead to G1 arrest in cells, however those cells which have low CDKN1A expression are halted in S phase in an ATR dependent manner. Cells paused in S phase either die through inability to restart fork progression in a SLFN11 dependent manner, or the DNA damage is repaired through homologous recombination. ATR inhibition with berzosertib treatment leads to a loss of the intra-S phase checkpoint and thus cells enter mitosis with unrepaired DNA damage and ultimately undergo mitotic catastrophe. XPG and p21 are determinants of response even in the presence of berzosertib as they are required for the initial DNA damage or for G1 arrest respectively, processes which are upstream of ATR dependent S phase arrest.

## Discussion

SCLC is a recalcitrant disease, and the majority of patients die of chemotherapy resistant disease. We determined that the ATR inhibitor berzosertib strongly synergized with the recently approved second line chemotherapeutic lurbinectedin in SCLC, particularly in MYC-high, chemo-resistant, non-NE models. Combination synergy is dependent on the ability of lurbinectedin to induce DNA damage, and berzosertib to augment this damage by inhibiting cell cycle checkpoints leading to mitotic catastrophe. We identified two main factors which determine response to the combination. First, the ability of lurbinectedin to cause DNA damage is reliant upon NER, and in particular the expression of *ERCC5*/XPG. Secondly, the expression of *CDKN1A*/p21 is predictive of synergy, with higher *CDKN1A*/p21 expression leading to decreased synergy. Importantly both factors are associated with NE differentiation and MYC overexpression, with recalcitrant high-MYC non-NE tumors displaying increased *ERCC5*/XPG and decreased *CDKN1A*/p21 leading to increased synergy of the combination.

SCLC may be particularly responsive to this combination due to the frequent mutations of *TP53* and thus decreased ability of cells to activate *CDKN1A*/p21 in response to DNA damage. Previous work has indicated that ATM inhibitors were required for ATR inhibitors to synergize effectively with lurbinectedin [50]. We found that in the majority of SCLC cell lines ATR and lurbinectedin synergized even in the absence of ATM inhibitors, potentially due to the high frequency of loss-of-function *TP53* mutations. Further, ATM inhibition was more impactful in the rare *TP53* proficient cells as compared to *TP53* mutant cells.

HR deficiency led to an increase in lurbinectedin efficacy. This observation is consistent with previous findings in breast cancer where HR-deficient breast cancers, particularly *BRCA2* mutant tumors, were more likely to respond to lurbinectedin than *BRCA1*/2 wild-type tumors [51]. Classical HR deficiency due to HR pathway member mutation (i.e. *BRCA1/2*) is rare in SCLC [9, 24-26]. However, based on our results in the BRCA2-KO DT40 cells where synergy was reduced but not abolished, we believe that the addition of berzosertib would still be meaningful in an HR deficient setting. This is supported by our results in the SCLC cell line NCI-H1048 in which lurbinectedin and berzosertib were synergistic. This is noteworthy as NCI-H1048 cells were ∼10 fold more sensitive to DNA damaging agents (lurbinectedin, topotecan) as compared to other SCLC cell lines tested and displayed low *BRCA2* expression indicating this line may be HR deficient or have reduced HR competency.

Consistent with recently published work [37], SLFN11 expression trended towards predicting lurbinectedin efficacy, and *SLFN11*-KO reduced lurbinectedin efficacy and increased lurbinectedin-berzosertib synergy. However, we found that SLFN11 expression across cell lines did not correlate with synergy, and that there was still significant synergy *in-vitro* and *in-vivo* even in SCLC models with high expression of SLFN11. While cell lines/tumors with high SLFN11 are more sensitive to lurbinectedin alone, thus reducing the impact of adding an ATR inhibitor, lurbinectedin-berzosertib synergy is not mechanistically dependent on SLFN11 expression levels. This is supported by *ERCC5*/XPG knockout cells which are also resistant to lurbinectedin, but unlike SLFN11 knockout cells they displayed decreased synergy and lurbinectedin efficacy cannot be rescued with the addition of berzosertib. Based on these data and the high degree of heterogeneity in second line SCLC we are not currently stratifying patients based on SLFN11 or HR status although we will continue to observe these parameters.

In a previous clinical trial, lurbinectedin in combination with the topoisomerase II inhibitor doxorubicin failed to improve survival as compared to standard of care [30]. This may be due to similar mechanisms of action for both agents as both lurbinectedin and doxorubicin cause DNA damage through poisoning DNA-protein interactions (lurbinectedin binds DNA and XPG while doxorubicin binds DNA and Topoisomerase II), likely leading to similar resistance and repair mechanisms (i.e. HR, SLFN11). We have demonstrated that lurbinectedin has much greater synergistic potential with ATR inhibitors, in particular berzosertib, than with doxorubicin as part of our initial screen. In light of the improved synergy of lurbinectedin and berzosertib as compared to doxorubicin, and the diverse mechanisms of action we believe lurbinectedin and berzosertib may have improved efficacy in SCLC patient as compared to lurbinectedin and doxorubicin.

Berzosertib is currently being assessed in combination with topotecan for the treatment of SCLC and has so far shown great promise leading to durable responses in several patients [5, 6]. Both topotecan and lurbinectedin synergized highly with berzosertib based on our work, and although lurbinectedin was more potent in 7/9 SCLC cell lines tested, due to different dosing schemes and bioavailability whether this will impact efficacy in patients is unclear. The combination of berzosertib and topotecan has however shown the greatest response in patients with high NE differentiation [6], whereas our work has shown that the lurbinectedin-berzosertib combination is likely to have the greatest degree of improvement in non-NE patients. In particular the increased *ERCC5*/XPG in non-NE tumors should only effect lurbinectedin-berzosertib combination efficacy, with little to no effect on topotecan-berzosertib combination efficacy. With continued assessment of biomarkers for both combinations, in future second line patients with NE SCLC could be treated with topotecan-berzosertib while patients with non-NE SCLC may be treated with lurbinectedin-berzosertib.

Based on these results, a phase I/II clinical trial with a combination of lurbinectedin and berzosertib was launched (NCT04802174) to determine anti-tumor efficacy in SCLC patients. In light of our findings described here, it will be important to assess whether NE differentiation along with *ERCC5*/XPG, *CDKN1A*/p21 and MYC family member expression are predictors for the efficacy of the combination treatment in patients.

## ACKNOWLEDGMENTS

This work was supported in part by the intramural programs of the Center for Cancer Research, National Cancer Institute (ZIA BC 011793) and EMD Serono (CrossRef Funder ID: 10.13039/100004755).

